# *iModMix*: Integrative Module Analysis for Multi-omics Data

**DOI:** 10.1101/2024.11.12.623208

**Authors:** Isis Narváez-Bandera, Ashley Lui, Yonatan Ayalew Mekonnen, Vanessa Rubio, Noah Sulman, Christopher Wilson, Hayley D. Ackerman, Oscar E. Ospina, Guillermo Gonzalez-Calderon, Elsa Flores, Qian Li, Ann Chen, Brooke Fridley, Paul Stewart

**Affiliations:** Department of Biostatistics and Bioinformatics, Moffitt Cancer Center; Department of Molecular Oncology, Moffitt Cancer Center; Cancer Biology and Evolution Program, Moffitt Cancer Center; Health Informatics Institute, University of South Florida; Department of Biostatistics, St. Jude Children’s Research Hospital; Huntsman Cancer Institute, University of Utah; Division of Health Services and Outcome Research, Children’s Mercy Research Institute; Department of Nutrition and Integrative Physiology, University of Utah

## Abstract

**Summary:** Integrative Module Analysis for Multi-omics Data (*iModMix*) is a biology-agnostic framework that enables the discovery of novel associations across any type of quantitative abundance data, including but not limited to transcriptomics, proteomics, and metabolomics. Instead of relying on pathway annotations or prior biological knowledge, *iModMix* constructs data-driven modules using graphical lasso to estimate sparse networks from omics features. These modules are summarized into eigenfeatures and correlated across datasets for horizontal integration, while preserving the distinct feature sets and interpretability of each omics type. *iModMix* operates directly on matrices containing expression or abundances for a wide range of features, including but not limited to genes, proteins, and metabolites. Because it does not rely on annotations (*e*.*g*., KEGG identifiers), it can seamlessly incorporate both identified and unidentified metabolites, addressing a key limitation of many existing metabolomics tools. *iModMix* is available as a user-friendly R Shiny application requiring no programming expertise (https://imodmix.moffitt.org), and as an R package for advanced users (https://github.com/biodatalab/iModMix). The tool includes several public and in-house datasets to illustrate its utility in identifying novel multi-omics relationships in diverse biological contexts.

**Availability and implementation:** Shiny application: https://imodmix.moffitt.org. The R package and source code: https://github.com/biodatalab/iModMix.

## Introduction

Integration of multiple omics modalities (i.e., “multi-omics”) is essential for a comprehensive understanding of complex biological systems and disease mechanisms. The scale and complexity of multi-omics data necessitate computational approaches that are adaptable to diverse experimental designs and data types, while remaining accessible to users without extensive programming experience. A key challenge in this space is the integration of untargeted metabolomics data, where a large fraction of detected metabolites are unidentified (Jendoubi, 2021). Current pathway analysis tools generally require standardized identifiers to map features to known biological pathways, and therefore they inherently cannot incorporate unidentified metabolites. Even for identified compounds, inconsistent naming conventions and incomplete annotation (e.g., missing KEGG identifiers) often prevent their use in pathway-based analyses. As a result, substantial portions of metabolomics data are excluded, limiting downstream analysis and reducing the potential for functional interpretation of results (Jendoubi, 2021).

In general, approaches for integrating multi-omics can be categorized into vertical or horizontal integration. Vertical integration takes multiple omics datasets as input and produces a single output, such as clustering assignments or a merged dataset, which no longer reflects the individual features of the original data. iClusterPlus (Qianxing et al., 2013) is an example of vertical integration: it takes multiple omics datasets from the same samples as input, extracts latent factors across omics datasets, clusters samples in latent space, and outputs the clustering assignments by sample. Another example of vertical integration is Non-negative Matrix Factorization (NMF) (Brunet et al., 2004) which is incorporated into tools like intNMF (Chalise & Fridley, 2017) which uses integrative NMF to cluster samples based on multiple genomic datasets. In contrast, horizontal integration analyzes multiple omics datasets in parallel to maintain the context of the original inputs. PIUmet (Pirhaji et al., 2016) is an example of horizontal integration: it takes metabolomics and proteomics data as input, and the resulting network output contains both metabolites and proteins.

*iModMix* is a horizontal integration framework that constructs network modules from any combination of input omics datasets (*e*.*g*., transcriptomics, proteomics, metabolomics). It first uses graphical lasso to estimate a sparse Gaussian Graphical Model (GGM) (Friedman et al., 2008) for each input dataset. GGMs capture direct associations within the input omics datasets, in constrast to Weighted Gene Correlation Network Analysis (WGCNA) (Langfelder, Horvath, et al., 2008), which is based on pairwise correlations and therefore includes both direct and indirect associations. Next, a Topological Overlap Matrix (TOM) is calculated (Yip et al., 2007) to quantify the extent to which pairs of features share common neighbors. Hierarchical clustering is then performed on TOM dissimilarity to group related features into modules. *iModMix* then finds the first principal component (PC) of the abundances of the features in the module, called an eigenfeature. These eigenfeatures can be used as abundance measurements for each module, and therefore can be used for testing for differential expression or association with experimental conditions at the module level. Finally, integration between omics types is achieved by correlating the eigenfeatures from different omics sets.

A major advantage of *iModMix* is its ability to generate network modules and associated eigenfeatures independent of feature annotation. In the case of metabolomics, *iModMix* incorporates both identified and unidentified metabolites into the analysis, overcoming a common limitation of many existing metabolomics tools. Although we emphasize metabolomics here, this same approach can be applied to other omics types where the identity or function of some features are unknown, such as long non-coding RNAs in transcriptomics. Because *iModMix* does not rely on existing pathway databases like KEGG, it also supports the discovery of novel associations between features. To highlight its functionality, we have applied *iModMix* to two case studies (see supplementary file). The first uses public data on clear cell renal cell carcinoma (ccRCC, RC20 dataset) to validate the results (Case Study S1) while the second demonstrates the utility of *iModMix* in analyzing untargeted metabolomics data that includes unidentified features (Case Study S2, Table S1, Fig. S1).

## Implementation

### Data Pre-Processing

Users can integrate any number of omics datasets simultaneously with the *iModMix* R package, while the Shiny application (https://imodmix.moffitt.org) supports up to three omics datasets (Fig. 1A). A separate metadata file containing sample IDs and at least one sample grouping column (*e*.*g*., treatment, control) is necessary for conducting PCA, heatmaps, and boxplots. Users can provide feature-level annotation corresponding to the input omics dataset. *iModMix* calculates the missing values in each row and filters out rows with missing values in more than 10% of the columns. It then calculates the standard deviation for each column and filters out columns with a standard deviation below the 25th percentile. Missing values are imputed using the k-nearest neighbors’ algorithm, or users can also provide their own imputed data if a different method is preferred. Finally, the data is scaled by centering each feature to have a mean of zero and a standard deviation of one. In addition to the case studies, we provide ten multi-omic datasets (transcriptomics and metabolomics) in the Shiny application to further illustrate *iModMix’s* capabilities. These datasets (Benedetti et al., 2023) have been formatted to meet *iModMix* specifications, with variable columns labeled as “Feature_ID” and sample columns labeled as “Samples”. These datasets are readily accessible for download on Zenodo (https://doi.org/10.5281/zenodo.13988161). Furthermore, we provide an in-house dataset of 20 lung adenocarcinoma (LUAD) mouse samples (10 wild type, 10 knockout), with 7353 metabolomics features and 7928 protein groups, to highlight *iModMix’s* functionality in handling unidentified metabolites. Analysis of these example datasets are described in more detail in the supplementary file.

**Figure 1.**
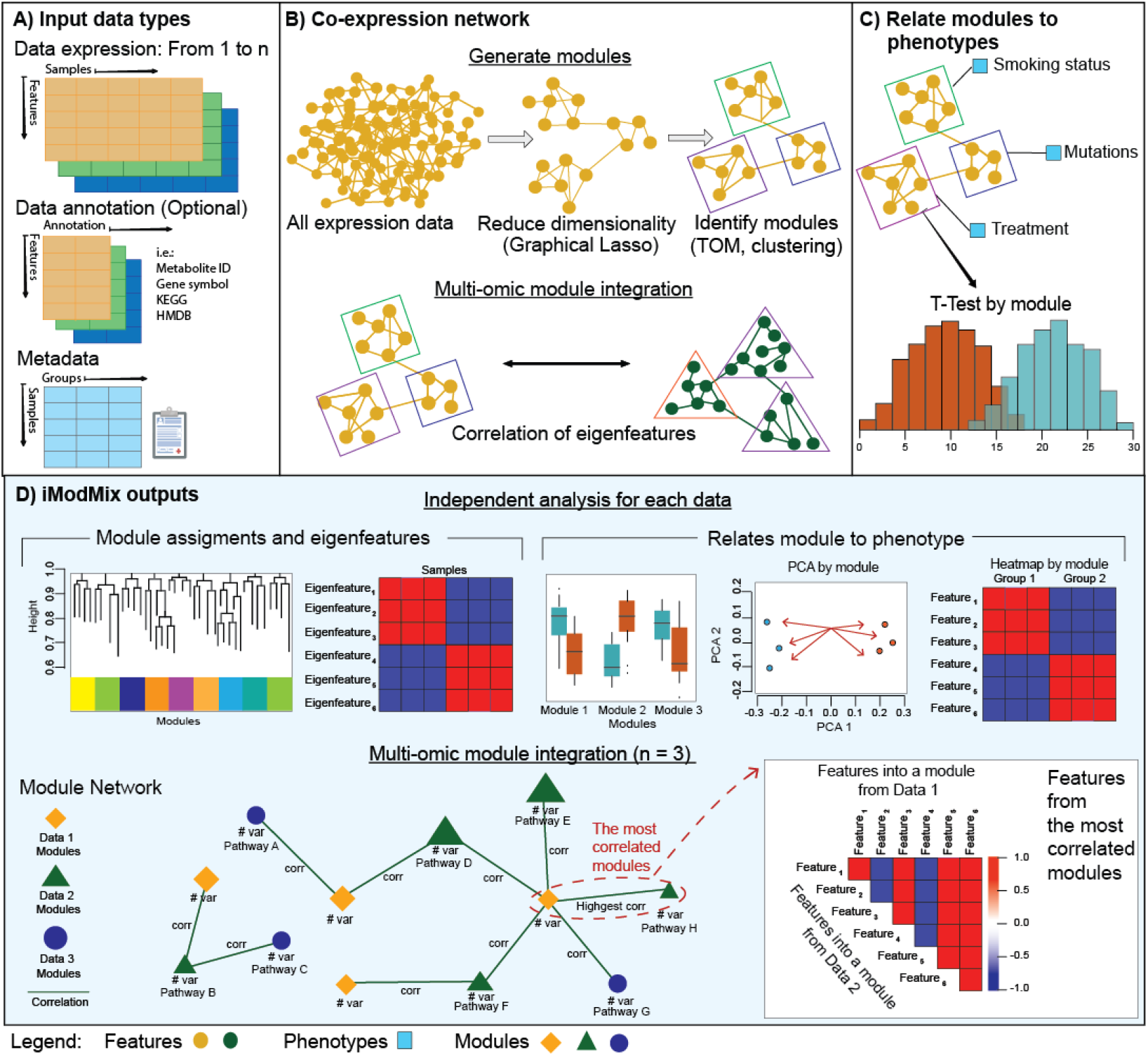
Overview of *iModMix*. (A) *iModMix* integrates multi-omics and clinical data to understand biological systems comprehensively. (B) Co-expression network: The tool performs independent analyses of each omics dataset to identify biologically relevant modules, and it then integrates the datasets by correlating eigenfeatures. (C) *iModMix* evaluates associations between phenotype-related modules across different data types. (D) Module assignments and eigenfeatures for each data set are calculated and phenotypes are compared. A module network detailing interactions between all datasets is generated along with the correlation plot for visualization.

### Module Construction

*iModMix* first estimates partial correlations using the glassoFast R package (Friedman et al., 2008) (Fig. 1B) to build a GGM. The regularization parameter lambda was set to a default value of 0.25 after benchmarking (see benchmarking section). The function also allows users to input their desired value for lambda using the *iModMix* R package. Next, the TOM is computed and TOM dissimilarity (1 – TOM) is used for hierarchical clustering (Yip et al., 2007). Modules are created using the dynamic tree cut method, specifically the ‘hybrid’ method, which refines assignments derived from a static cut height by analyzing the shape of each branch (Langfelder, Zhang, et al., 2008). Finally, *iModMix* calculates eigenfeatures for each module (Fig. 1B) for downstream omics analyses.

### Module Exploration

The phenotype analysis section provides a framework for classifying phenotypes based on eigenfeatures. Users can upload metadata, select phenotypes of interest, and set significance thresholds for statistical tests. Significant eigenfeatures are visualized through boxplots. Additionally, users can explore specific modules and view associated features, PCA loading, and heatmap plots to view feature behavior across different phenotypes (Fig. 1C). *iModMix* supports pathway enrichment analysis of genes or proteins by utilizing all available libraries in Enrichr if gene symbols are provided (Kuleshov et al., 2016).

### Multi-omics analysis

*iModMix* integrates modules across all available omics datasets by calculating the Spearman correlation between eigenfeatures for all possible combinations. This is followed by construction of an interactive network, where modules from Data 1 are represented as yellow diamonds, modules from Data 2 as green triangles, modules from Data 3 as blue circles, and correlation coefficients on connecting arrows. Users can select the number of top multi-omics module correlations to visualize and explore detailed correlations within these modules. Lists of metabolites and proteins/genes within highly correlated modules are provided, facilitating pathway analysis and offering insights into the relationships between different omics layers. Additionally, classification between features of each layer and phenotypes using t-tests and boxplots is provided, enabling users to visualize and identify differences (Fig. 1B).

### System requirements

*iModMix* can run on standard computers with > 8GB of RAM and a multi-core processor. The running time depends on data size and complexity. Typical data sizes of 20,000 variables can be completed in around 10 minutes (Fig. S2).

### Benchmarking

In *iModMix*, the regularization parameter lambda (λ) controls the sparsity of the partial correlation matrix estimated using the glasso package (Friedman et al., 2008). This matrix determines which direct associations between features are retained and which are removed. Because the network structure derived from these partial correlations forms the basis for downstream module detection, λ is a critical parameter. If λ is too high, the resulting network is overly sparse and may not capture meaningful relationships. If λ is too low, the network retains too many weak associations, which can lead to large, uninformative modules or overly fragmented module structures, depending on the clustering method used. To identify a practical default, we benchmarked *iModMix* on datasets from clear cell renal cell carcinoma (ccRCC, RC20 dataset) (Benedetti et al., 2023), and lung adenocarcinoma. For each dataset, we applied the *iModMix* pipeline and got a λ = 0.25 for consistently balanced speed and module quality (Fig. S3). This value is set as the default in *iModMix* Shiny app. Advanced users can modify this parameter in the R package to suit their data.

*iModMix* was also compared directly with WGCNA using the lung adenocarcinoma (LUAD) data. For metabolomics data, WGCNA generated 9 modules (Mean size = 817, SD = 2206), whereas *iModMix* produced 287 modules (Mean size = 23, SD = 11.2). For proteomics data, WGCNA generated 22 modules (Mean size = 360, SD = 559), while *iModMix* identified 412 modules (Mean size = 17, SD = 5.75) (Table S2, Fig. S4). WGCNA yielded a wide range of module sizes with one or two large modules comprising most features, whereas *iModMix* modules were more uniform in size (Fig. S3, Fig. S5). Smaller module sizes generated from *iModMix* resulted in more comprehensible grouping for pathway enrichment, identifying 181 pathways from proteins modules including all 13 pathways identified by WGCNA (Fig. S6, Table S3). The correlation in the integration process between metabolite and protein modules was higher with *iModMix* (0.93) compared to WGCNA (0.88) (Fig. S7). This benchmarking process demonstrates that *iModMix* generates more evenly sized modules with less features per group by using only direct associations for network construction, thus allowing easier interpretation and downstream analysis. A comparative overview of the underlying principles and analytical frameworks of *iModMix*, PIUMet, iClusterPlus, and WGCNA is presented in the Table S4.

### Summary

The *iModMix* pipeline provides an empirical framework for uncovering novel associations between multi-omics data, with the unique capability to incorporate both identified and unidentified metabolites. By generating *de novo* network modules independent of pathway databases, *iModMix* enables the discovery of previously unrecognized interactions between features. Unlike methods that rely on predefined biological functions or annotations, *iModMix* leverages feature matrices alone, making it highly adaptable to a wide range of omics combinations, including those with limited annotation or poorly characterized interactions.

