## Supplementary file 1. for "*iModMix*: Integrative Module Analysis for Multi-omics Data"

**Case Study S1. Clear cell renal cell carcinoma (ccRCC, RC20) Dataset with Identified Metabolites**

We used 24 normal and 52 tumor clear cell renal cell carcinoma (ccRCC) samples (Golkaram et al. 2022; Tang et al. 2023; Benedetti et al. 2023) as a case study. It contained 23001 genes from RNA-seq and 904 identified metabolites from untargeted metabolomics. Applying iModMix identified 751 gene modules and 34 metabolite modules. Differential expression analysis of the modules through t-test confirmed changes in metabolite abundance between groups recapitulated those from prior work (Benedetti et al. 2023) highlighting reduced levels of gamma-glutamyltyrosine (module ME#BA3241, P-value: 0.0003), creatinine/C00791 (module ME#C06162 P-value: 4.2681E-15), xanthosine/C01762 (module ME#66628D P-value: 0.0015), docosahexaenoate (DHA; 22:6n3)/C06429 (module ME#904A67 P-value: 4.6906E-12), and 1-methyladenosine/C02494 (module ME#E1C62F P-value: 8.1847E-14) in tumors compared to normal tissue. Conversely, metabolites with increased abundance in tumors compared to normal tissues included proline/C00148, glutamine/C00064 (module ME#6C856F, P-value: 0.006), maltose/C00208 and 1-methylnicotinamide/C02918 (module ME#904A67 P-value: 4.6906E-12) (Benedetti et al. 2023). The top five most correlated metabolomic and transcriptomic module pairs included pathways such as Inflammatory Response (P-value: 0.0001), Protein Polyubiquitination (P-value: 0.0001), tRNA Aminoacylation for Mitochondrial Protein Translation (P-value: 0.0084), and Sphingosine-1-Phosphate Receptor Signaling Pathway (P-value: 0.0068). The inflammatory response plays a crucial role in the development and prognosis of ccRCC. Research indicates that inflammation is involved at all stages of the disease, influencing tumor progression and treatment responses (Zhong et al. 2023).

**Case Study S2. Lung Adenocarcinoma Dataset with Unidentified Metabolites**

Matched proteomics and metabolomics dataset was generated using two mouse models for lung adenocarcinoma (LUAD) (10 wild type, 10 knockout), with 7353 metabolomics features (identified and unidentified) and 7928 protein groups. Applying iModMix generated 412 gene modules, and 287 metabolite modules. Strong correlations were observed between these modules, with correlations as high as 0.93. The correlations observed between variables in the 2 and 3 correlated pairs (Table S1) are notably high, with numerous correlations between proteins and both identified and unidentified metabolites exceeding 0.9 (Fig. S1A, Fig. S1B). These strong correlations offer valuable insights into the unidentified metabolites, providing crucial information about their potential relationships and helping to elucidate their behavior. Pathway analysis of these 16 genes from and the identified metabolite using MetaboAnalyst (Pang et al. 2024) revealed significant enrichment in Proteoglycans in cancer (P-value: 7.9492E-4) and Renin-angiotensin system (P-value: 7.9861E-4). Proteoglycans are critical in LUAD, influencing tumor progression, metastasis, and microenvironment. Their dysregulation impacts essential cellular processes such as epithelial-to-mesenchymal transition and cancer stemness, which are key to cancer cell plasticity and aggressiveness (Karagiorgou et al. 2022). Besides, focal adhesions play a significant role in LUAD progression by mediating cell adhesion, migration, and signaling pathways essential for tumorigenesis. Studies highlight the involvement of focal adhesion kinase (FAK) in LUAD, suggesting potential therapeutic targets and biomarkers for this cancer type (Zhou et al. 2018).

**Supplementary Table 1. Top 3 modules Case Study 2:** Results from iModMix analysis of Lung Adenocarcinoma (LUAD) dataset with unidentified metabolites highlighting the top correlated metabolomic-proteomic modules.

| Rank | Metabolite Module | # of var | Metabolites | Protein Module | # of var | Proteins | Corr |
|---|---|---|---|---|---|---|---|
| 1 | #BF862B | 18 | 0 identified | #CF8BAF | 11 | Pdgfra, Klhl15, Stx2, Rtkn2, Ager, Gypc, Erbin, Rap2a, Abca8b, Lims2, Cav2 | 0.9293 |
| 2 | #FFB214 | 21 | 7 identified:<br>C00166: Phenylpyruvic acid,<br>C00954: Indoleacetic acid,<br>C05582: Homovanillic,<br>C03672: Hydroxyphenyllactic acid,<br>C01179: 4Hydroxyphenylpyruvic,<br>C00526: Deoxyuridine,<br>C00178: Thymine | #FFDD25 | 16 | Myl6, Tmod1, Vsnl1, Tgfb1i1, Ppp1r14a, Aldh1a1, Ilk, Myl12b, Dlc1, Plscr4, Macf1, Pcdh18, Limch1, Lims1, Specc1l, Vcl, Rasip1 | 0.9293 |
| 3 | #5E985E | 26 | 1 identified:<br>C06426: gamma-linolenic acid | #FFDA24 | 16 | Hsd11b1, Cavin2, Ehd2, Myh10, Clic5, Msn, Myo1c, Epb41l2, Pakap, Sptan1, Akap12, Cyp2b10, Cav1, Cav3, Sptbn1, Ace, Ace3, Cavin1 | 0.9278 |

**Supplementary Figure 1. Top modules Case Study 2:** Results from *iModMix* analysis of Lung Adenocarcinoma (LUAD) dataset using metabolomics with unidentified metabolites. Results highlight the top correlated metabolomic-proteomic modules with unidentified metabolites.

A) Most correlated modules: Top 2

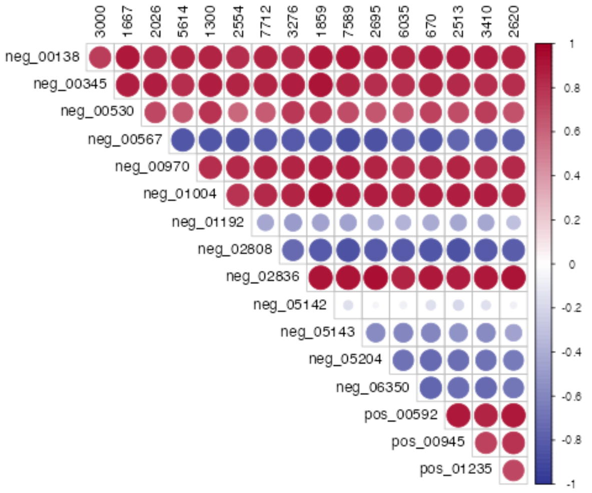


Modules correlation: Data 1 and Data 2

Show 10 entries    Search:

| | Data1 | Data2 | Correlation |
|---|---|---|---|
| 203 | pos_02119 | 699 | 0.9414 |
| 204 | pos_02244 | 699 | 0.9414 |
| 369 | neg_00999 | 1023 | 0.9338 |
| 126 | pos_02244 | 4567 | 0.9278 |
| 334 | pos_02244 | 1668 | 0.9248 |
| 48 | pos_02244 | 3331 | 0.9233 |
| 125 | pos_02119 | 4567 | 0.9233 |
| 308 | pos_02244 | 1103 | 0.9218 |
| 411 | pos_02119 | 598 | 0.9218 |
| 412 | pos_02244 | 598 | 0.9218 |

Showing 1 to 10 of 416 entries    Previous 1 2 3 4 5 … 42 Next

Features from Data 1

| | Feature_ID | Metabolite |
|---|---|---|
| 4 | neg_00567 | Thymine |
| 10 | neg_05142 | Phenylpyruvate |
| 13 | neg_06350 | Indole-3-acetate |
| 7 | neg_01192 | Homovanillate\|3-(4-Hydroxyphenyl)lactate |
| 8 | neg_02808 | Deoxyuridine |
| 11 | neg_05143 | 3-(4-Hydroxyphenyl)pyruvate |

Features from Data 2

| | Feature_ID | Symbol |
|---|---|---|
| 1 | 1300 | Aldh1a1 |
| 2 | 1667 | Tmod1 |
| 3 | 1859 | Plscr4 |
| 4 | 2026 | Vsnl1 |
| 5 | 2513 | Speccl1 |
| 6 | 2554 | Myl12b |
| 7 | 2620 | Rasip1 |
| 8 | 2695 | Limch1 |
| 9 | 3000 | Myl6 |
| 10 | 3276 | Tgfb1i1 |

| | Feature_ID | Symbol |
|---|---|---|
| 11 | 3410 | Vcl |
| 12 | 5614 | Ppp1r14a |
| 13 | 6035 | Lims1 |
| 14 | 670 | Ilk |
| 15 | 7589 | Macf1 Pcdh18 |
| 16 | 7712 | Dlc1 |

B) Most correlated modules: Top 3

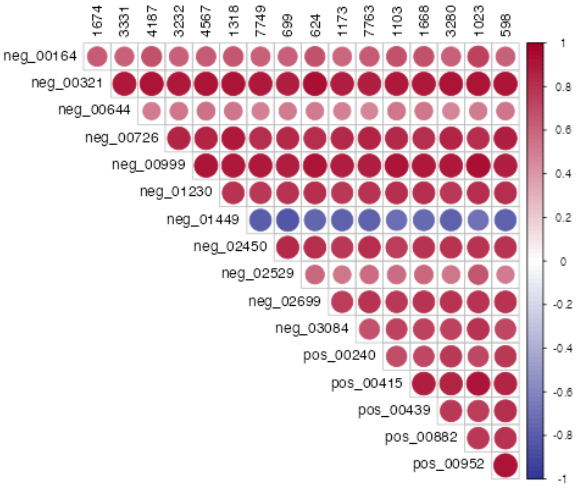


Modules correlation: Data 1 and Data 2

Show 10 entries    Search:

| | Data1 | Data2 | Correlation |
|---|---|---|---|
| 38 | pos_01316 | 1667 | -0.9338 |
| 185 | pos_01316 | 1859 | -0.9323 |
| 219 | neg_[illegible]836 | 2695 | 0.9218 |
| 41 | pos_02525 | 1667 | 0.9158 |
| 164 | pos_01316 | 3276 | -0.9158 |
| 30 | neg_02836 | 1667 | 0.9143 |
| 198 | neg_02836 | 7589 | 0.9128 |
| 101 | pos_01316 | 1300 | -0.9113 |
| 203 | pos_00592 | 7589 | 0.9113 |
| 269 | pos_01316 | 670 | -0.9113 |

Showing 1 to 10 of 336 entries    Previous 1 2 3 4 5 … 34 Next

Features from Data 1

| | Feature_ID | Metabolite |
|---|---|---|
| 17 | pos_01023 | Gamma-linolenic acid |

Features from Data 2

| | Feature_ID | Symbol |
|---|---|---|
| 1 | 1023 | Ace Ace3 |
| 2 | 1103 | Cyp2b10 |
| 3 | 1173 | Sptan1 |
| 4 | 1318 | Msn |
| 5 | 1668 | Cav1 Cav3 |
| 6 | 1674 | Hsd11b1 |
| 7 | 3232 | Myh10 |
| 8 | 3280 | Sptbn1 |
| 9 | 3331 | Cavin2 |
| 10 | 4187 | Ehd2 |

| | Feature_ID | Symbol |
|---|---|---|
| 11 | 4567 | Clic5 |
| 12 | 598 | Cavin1 |
| 13 | 624 | Pakap |
| 14 | 699 | Epb41l2 |
| 15 | 7749 | Myo1c |
| 16 | 7763 | Akap12 |

**Supplementary Figure 2. Running time of *iModMix*.** The running time depends on the size and complexity of the dataset. Here we show 3 datasets demonstrating an *iModMix* run can be completed in around 10 minutes: Data 1: 7323 variables and 20 samples; Data 2: 7928 and 20 samples; Data 3: 23000 variables and 77 samples. can be completed in around 10 minutes

Data 1: 7323 variables

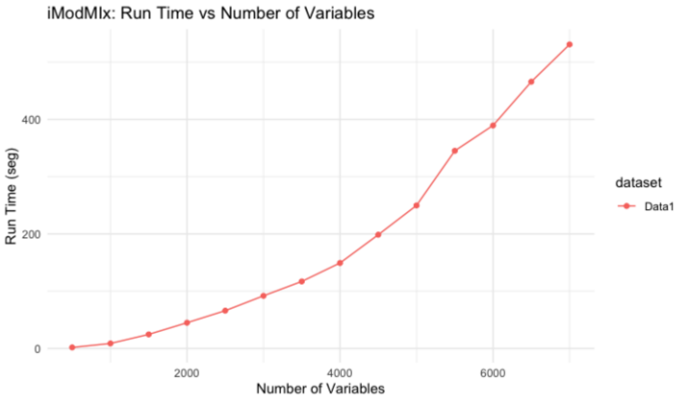


Data 2: 7928 variables

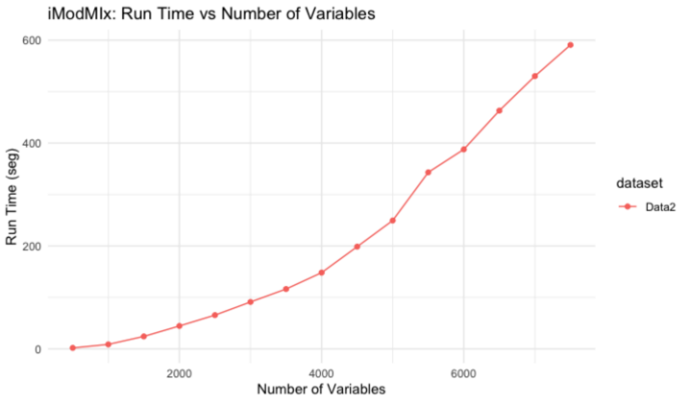


Data 3: 23000 variables

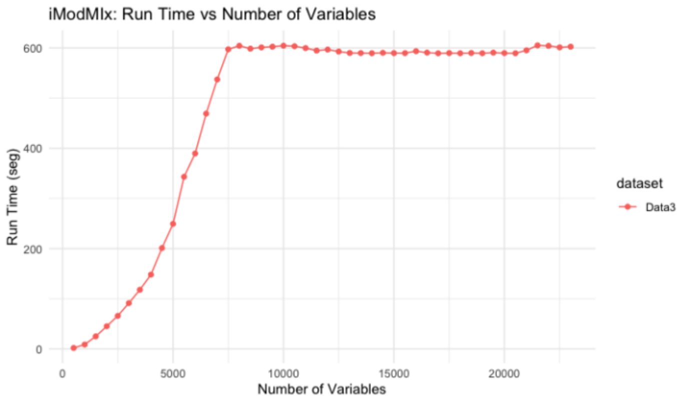


Data 1, Data 2 and Data 3

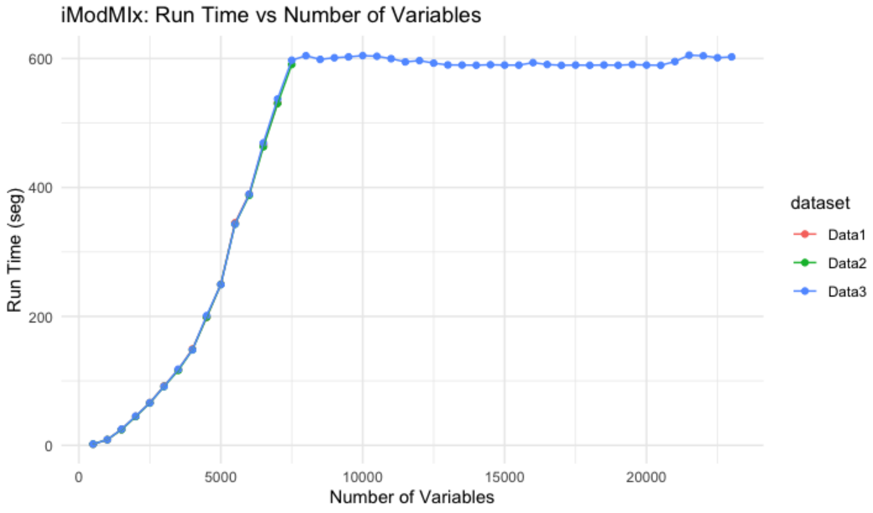

**Supplementary Figure 3. Benchmaking results.** Benchmarking using clear cell renal cell carcinoma pubic available data (ccRCC, RC20) and lung adenocarcinoma (LUAD) data results demonstrates the performance of lambda vs time for number of genes. Stability was achieved in both datasets with an alpha value of 0.25

A) Metabolomic abundance ccRCC (RC20 dataset)

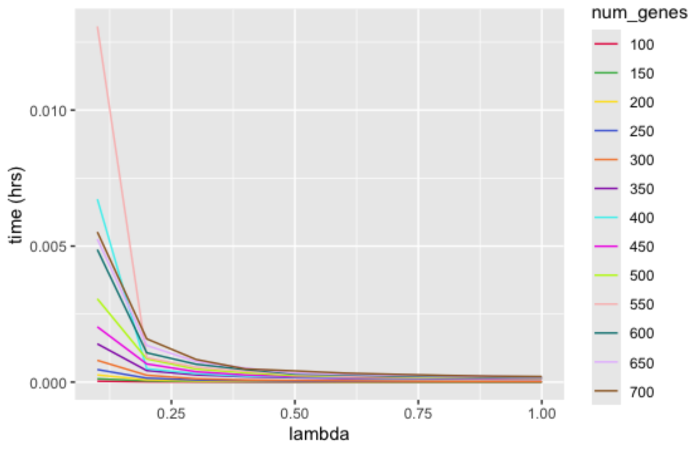


B) Gene expression lung adenocarcinoma ( LUAD) data

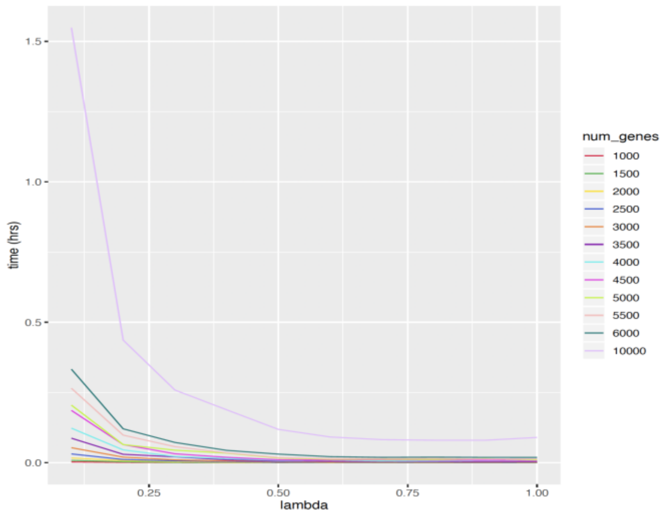


C) Gene expression lung adenocarcinoma ( LUAD) data

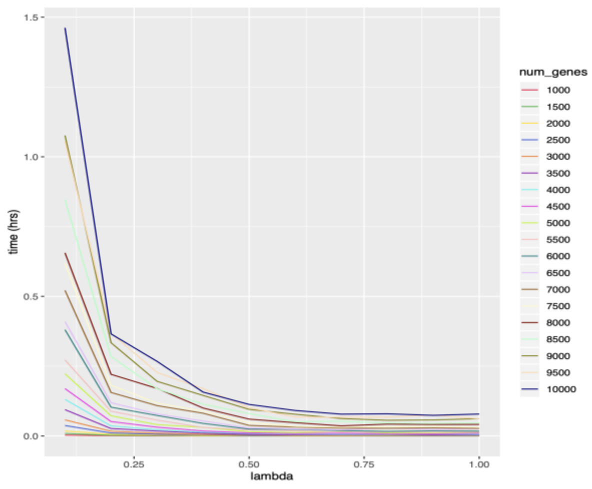

**Supplementary Table 2. WGCNA vs *iModMix*:** Number of modules generated from lung adenocarcinoma (LUAD) data. For metabolomics data, WGCNA generated 9 modules (Mean size = 817), whereas *iModMix* produced 287 modules (Mean size= 23). In the case of proteomics data, WGCNA generated 22 modules (Mean size= 360), while *iModMix* identified 412 modules (Mean size= 17). *iModMix* resulted in more modules with a smaller average number of features per module.

| Dataset | # of Features | WGCNA | | iModMix | |
|---|---|---|---|---|---|
| | | # Modules | Average of features per module | # Modules | Average of features per module |
| Metabolites | 7,353 | 9 | 817 | 287 | 23 |
| Proteins | 7,928 | 22 | 360 | 412 | 17 |

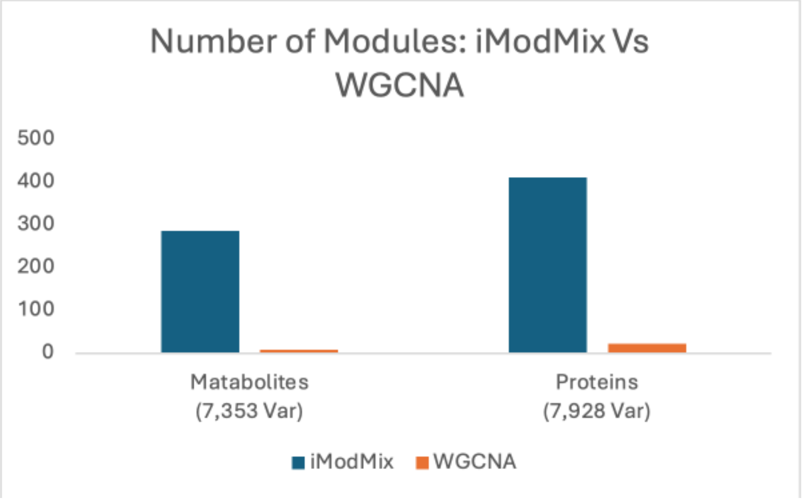

**Supplementary Figure 4. WGCNA vs *iModMix*:** Modules distribution in lung adenocarcinoma (LUAD) data shows *iModMix* resulted in smaller and more uniformly sized modules. For metabolomics data, WGCNA generated 9 modules (Mean size = 817, SD = 2206), whereas *iModMix* produced 287 modules (Mean size= 23, SD = 11.2). In the case of proteomics data, WGCNA generated 22 modules (Mean size= 360, SD = 559), while *iModMix* identified 412 modules (Mean size= 17, SD = 5.75).

Metabolomics LUAD data

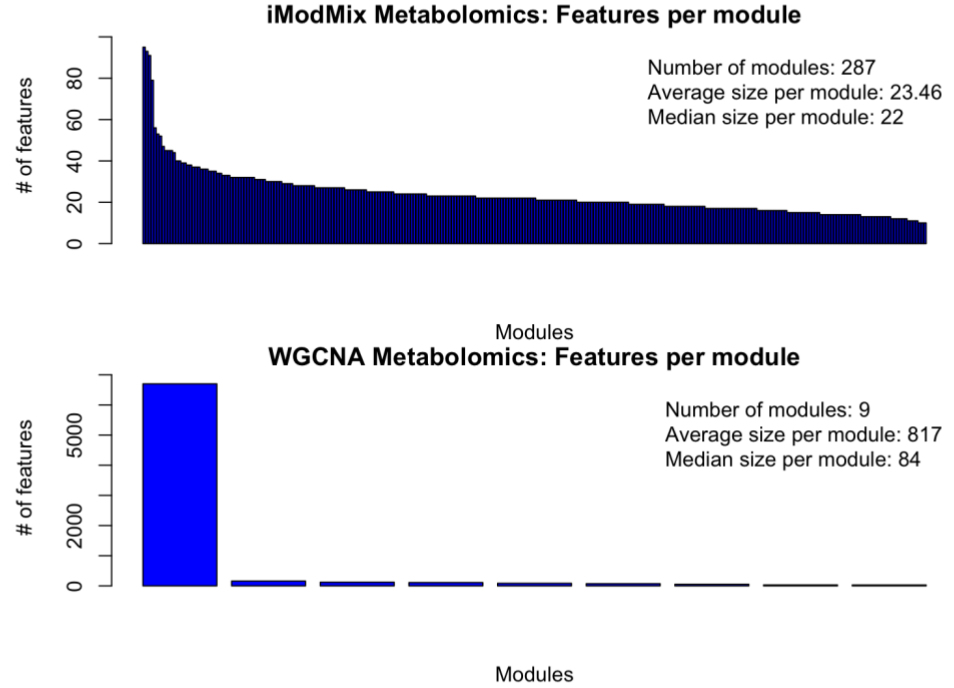


Proteomics LUAD Data

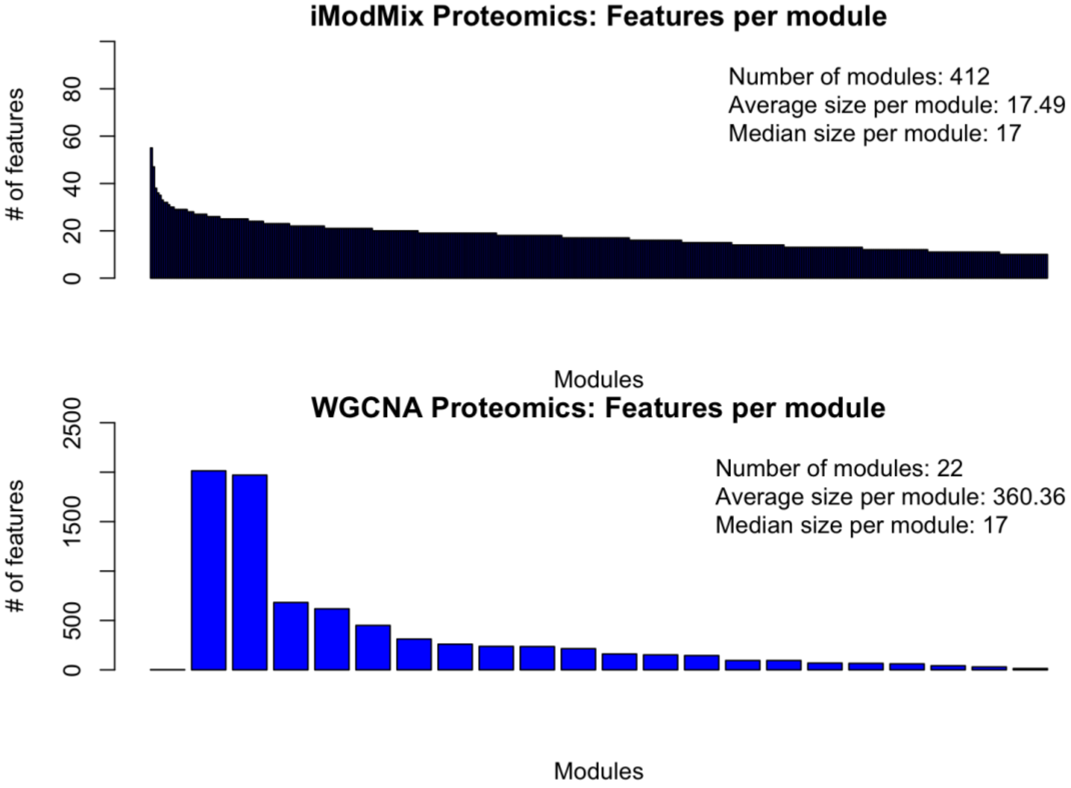

**Supplementary Figure 5. WGCNA vs *iModMix*:** Modules distribution in lung adenocarcinoma (LUAD) data. The standard deviation for WGCNA modules was very high, indicating a wide range of module sizes, whereas *iModMix* modules were more uniform in size (Fig. S3).

**Metabolomics data**

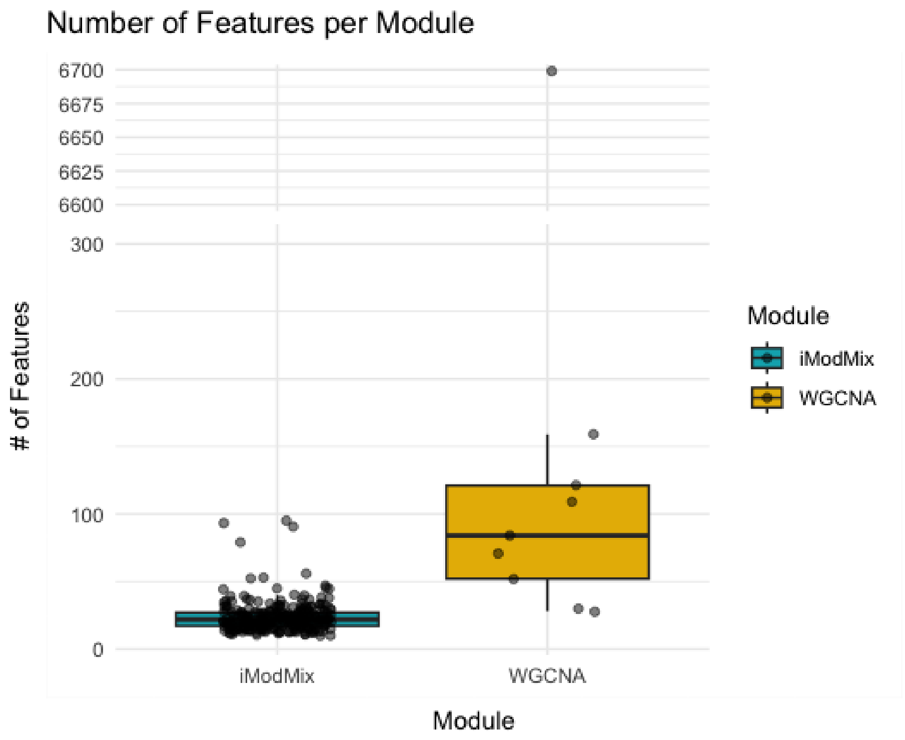


**Proteomics Data**

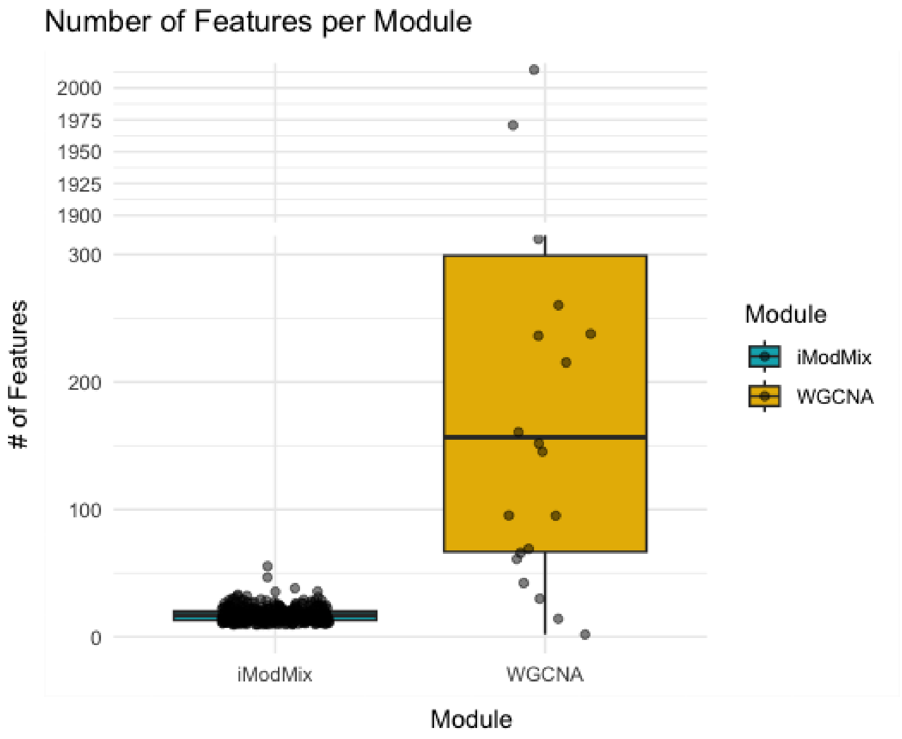

**Supplementary Figure 6. WGCNA vs *iModMix***: Enrichment Analysis by Enrichr for proteins modules using the lung adenocarcinoma (LUAD) data. *iModMix* resulted in smaller more uniformly sized modules, identifying 181 pathways (provided as a supplemental spreadsheet, Table S3) from protein modules compared to the 13 pathways identified by WGCNA. Notably, all 13 pathways identified by WGCNA were also detected by *iModMix*.

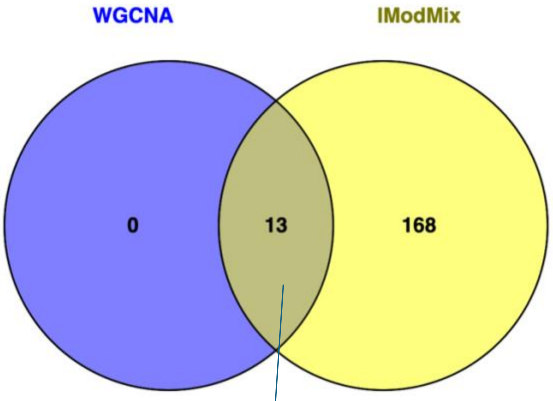


| **Commons** |
| --- |
| Ubiquitin mediated proteolysis |
| Platelet activation |
| Complement and coagulation cascades |
| Epstein-Barr virus infection |
| Valine, leucine and isoleucine degradation |
| Spliceosome |
| ECM-receptor interaction |
| Cardiac muscle contraction |
| Protein export |
| Oxidative phosphorylation |
| Metabolism of xenobiotics by cytochrome P450 |
| IL-17 signaling pathway |
| Ribosome |

**Supplementary Figure 7. Integration of omics data using WGCNA Vs *iModMix*.** Correlation plot from the integration process between metabolite and protein modules using the lung adenocarcinoma (LUAD) data. The top correlation achieved using *iModMix* was 0.93, whereas WGCNA reached a top correlation of 0.88.

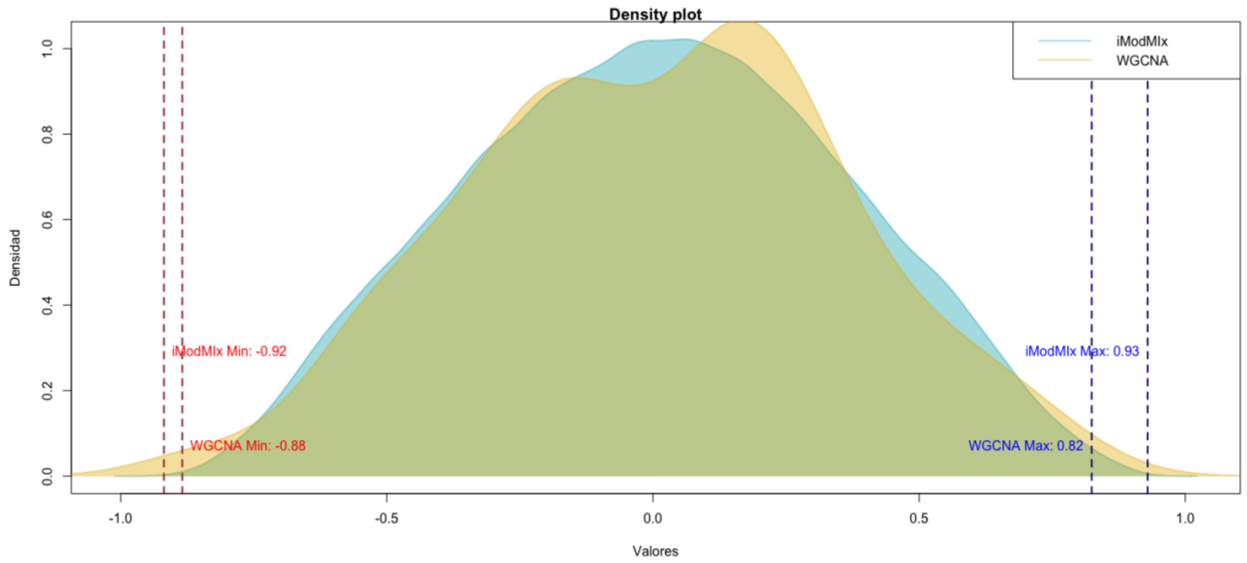


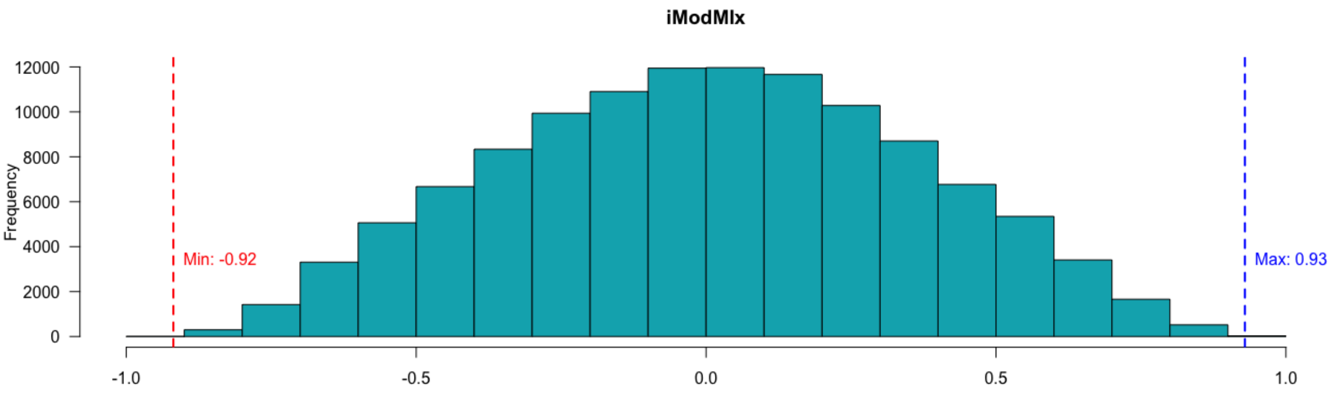


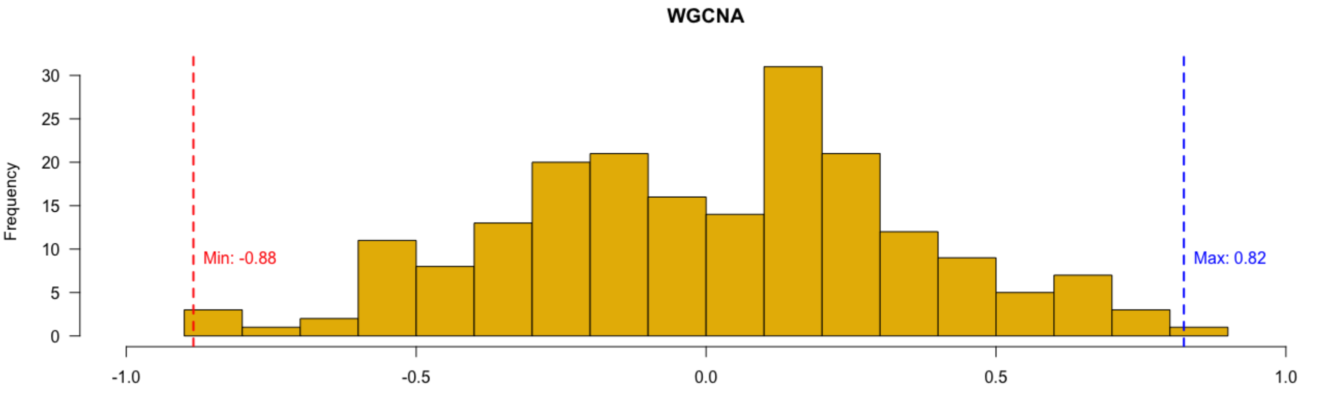


Top 5 correlations from integration

| Data1 | Data2 | Correlation |
|---|---|---|
| D1#red | D2#magenta | -0.8842 |
| D1#black | D2#magenta | -0.8496 |
| D1#turquoise | D2#turquoise | 0.8241 |
| D1#brown | D2#magenta | -0.8045 |
| D1#brown | D2#green | 0.7594 |

| Data1 | Data2 | Correlation |
|---|---|---|
| D1#BF862B | D2#CF8BAF | 0.9293 |
| D1#FFB214 | D2#FFDD25 | 0.9293 |
| D1#5E985E | D2#FFDA24 | 0.9278 |
| D1#B83343 | D2#DD1D22 | 0.9248 |
| D1#EF7718 | D2#DD1D22 | 0.9248 |

**Supplementary Table 4.** A comparative overview of the underlying principles and analytical frameworks of *iModMix*, PIUMet, iClusterPlus, and WGCNA.

| Feature / Method | *iModMix* | WGCNA | PIUMet | iClusterPlus |
|---|---|---|---|---|
| **Integration of multi-omics data** | Yes | No | Yes | Yes |
| **Supports Horizontal integration of multi-omics data** | Yes | Yes | Yes | No |
| **Processes any number of omics datasets simultaneously** | Yes | No | No | No |
| **Handling of unidentified metabolites** | Yes | No | Yes | No |
| **Employes network-based approach** | Yes | Yes | Yes | No |
| **Is not constrained by prior knowledge (e.g., seeds)** | Yes | Yes | No | Yes |
| **Identifies co-expression modules** | Yes | Yes | No | No |
| **Useful for feature selection** | Yes | Yes | Yes | Yes |
| **Provided as an R package** | Yes | Yes | No | Yes |
| **Allow to unveil novel connections** | Yes | Yes | Yes | Yes |
| **User-friendly (Shiny app)** | Yes | Yes | No | No |
| **Scores (Total Yes)** | 11 | 8 | 6 | 5 |
